# Vascular endothelial growth factor (VEGF) expression and neuroinflammation is increased in the frontopolar cortex of individuals with autism spectrum disorder

**DOI:** 10.1101/627083

**Authors:** Aswini Gnanasekaran, Megan N. Kelchen, Nicole K. Brogden, Ryan M. Smith

## Abstract

Autism spectrum disorder (ASD) etiology is a complex mixture of genetic and environmental factors, the relative contributions of which varies across patients. Despite complex etiology, researchers observe consistent neurodevelopmental features in ASD patients, notably atypical forebrain cortical development. Growth factors, cytokines, and chemokines are important mediators of forebrain cortical development, but have not been thoroughly examined in brain tissues from individuals with autism. Here, we performed an integrative analysis of RNA and protein expression using frontopolar cortex tissues dissected from individuals with ASD and controls, hypothesizing that ASD patients will exhibit aberrant expression of growth factors, cytokines, and chemokines critical for neurodevelopment. We performed group-wise comparisons of RNA expression via RNA-Seq and growth factor, cytokine, and chemokine expression via multiplex enzyme-linked immunosorbent assay (ELISA). We also analyzed single cell sequencing data from the frontopolar cortex of typically developed individuals to identify cell types that express the growth factors we found differentially expressed in ASD. Our RNA-Seq analysis revealed 11 differentially expressed genes in ASD versus control brains, the most significant of which encodes for vascular endothelial growth factor (VEGF-A). Both RNA and protein levels of VEGF-A were upregulated in ASD brains. Our single cell analysis revealed that VEGF is expressed primarily by non-neuronal cells. We also found that the differentially expressed genes from our RNA-Seq analysis are enriched in microglia. The increased VEGF-A expression we observed in ASD, coupled with the enrichment of differentially expressed genes in microglia, begs the question of the role VEGF-A is playing in ASD. Microglia activation, as indicated by our RNA-Seq results, and the VEGF-A isoform expression we see in the ASD cortex, leads us to conclude that VEGF-A is playing a pro-inflammatory role, perhaps with unwanted long-term consequences for neurodevelopment.

## Introduction

Autism spectrum disorders (ASD) constitute a variety of neurodevelopmental disorders defined clinically by difficulties in socialization, restricted or repetitive behaviors, and language delays, with childhood onset. ASD can be accompanied by concomitant symptoms including intellectual disability, catatonia, seizures, attention deficit hyperactivity disorder, and gastrointestinal dysfunction [1]. The cause of ASD is complex and involves both genetic and environmental risk factors.

The diagnostic clinical signs of ASD are preceded by many other phenotypic changes, including abnormal brain overgrowth in multiple brain regions. The frontal cortex, which is involved in higher-order social, emotional, communicative, and cognitive functions is implicated amongst these overgrowth regions [2,3] and exhibits abnormal laminar structure [4]. Functionally, the anterior frontal cortex (frontopolar cortex; Brodmann Area 10 – BA10) in individuals with ASD exhibits altered interhemispheric connectivity, which correlates with social deficits [5]. Others have found changes in temporal dynamics of resting state activity between the frontopolar cortex and other brain regions that correlate with attention deficits in ASD [6]. The abundance of evidence implicating the frontal cortex in ASD forms the basis of a unifying theory of ASD etiology [7], making this brain region an important focus for molecular and genetic analysis.

Genetic studies have started to unravel the molecular basis of autism. Genome wide associations [8] and whole-exome sequencing [9–11] reveal hundreds of rare and common inherited and *de novo* variants that contribute to autism risk. These variants often implicate processes related to neurodevelopment and neuronal function [8,12–14]. Neuroglial are critical for typical neural and synaptic development and modulating neuroinflammation, placing them central to the notable dysfunctions in ASD [15]. Genomic studies show that glial cell-mediated neuroinflammation is upregulated in the brains of individuals with ASD [16–18]. With respect to the frontal cortex, epigenetic changes in ASD occur adjacent to genes responsible for immune functions and synaptic maturation [19,20]. Increased pro-inflammatory factors are also found in peripheral blood [21,22] and brains from ASD patients [23]. Given these known relationships between neuroinflammation, neurodevelopment, and ASD, further understanding how these factors interact to underlie ASD etiology could lead to novel interventions that prevent or abate atypical neurodevelopment.

Several growth factors, secreted from both neurons and glial cells, play indispensable roles in shaping the brain [24]. VEGF-A promotes neuronal survival, axonal outgrowth, long-term potentiation, and learning [25,26]. However, VEGF-A signaling from reactive astrocytes can also drive vascular permeability and central nervous system (CNS) damage in acute inflammatory lesions [27]. Factors such as oxidative stress can result in enhanced glial expression of VEGF-A [28]. Serum, plasma, and cerebral spinal fluid (CSF) level of VEGF-A, VEGF receptors, and numerous other cytokines and chemokines are altered in patients with severe autism [29–32]. However, the significance of these finding when they occur in the periphery, as it relates to neurodevelopment, is unclear.

Given the apparent associations between cytokine and chemokine signaling, neurodevelopment, and ASD, our overall aim is to clarify the interactions between these factors using a comprehensive genomic approach in frontopolar cortex brain tissues from individuals with ASD and matched controls. Analysis of multiple chemokines and cytokines, whole transcriptome RNA-Seq profiling, and single nuclear RNA-Seq reveals convergent evidence that VEGF-A is upregulated in the cortex of ASD subjects, likely contributing to neuroinflammation.

## Materials and Methods

### Samples & RNA Isolation

Brain tissues were obtained through the Autism Tissue Program. They were dissected postmortem from the frontopolar cortex (BA10) of individuals with ASD (*n*=17) and controls *(n*=14). Subjects were predominately male (26M: 5F), white (28 white, 3 African-American), and ranged in age from 2-57 years old. We isolated RNA from these tissues using TRIzol (Invitrogen, Carlsbad, CA) and column purified the RNA with Qiagen RNeasy columns (QIAGEN, Germantown, MD), performing on-the-column DNase I treatment. RNA integrity was measured prior to RNA-Seq analysis and all samples had and RNA Integrity Number (RIN) > 6, indicating adequate quality for RNA expression measures. However, sample quality did decline, as determined by RNA integrity, following RNA-Seq analysis but prior to measuring cytokine and chemokine protein levels, which was accounted for in our statistical analyses. Table 1 lists demographics comparisons across the case versus control groups for each of the studies described below.

**Table 1.** Sample Demographics Across Experiments.

| Full Cohort |  |  |  |  |  |
| --- | --- | --- | --- | --- | --- |
| Diagnosis | Male:Female | Race | Age (S.D.) | RIN (S.D.) | PMI (S.D.) |
| ASD (n=21) | 16:5 | EA: 18, AA: 3 | 28.4 (20.7) | 7.6 (1.4) | **17.3 (6.0)\*** |
| Control (n=15) | 14:1 | EA: 14, AA: 1 | 34.3 (16.9) | 7.8 (1.0) | **21.9 (5.0)\*** |
| RNA-Seq |  |  |  |  |  |
| ASD (n=16) | 13:3 | EA: 14, AA: 2 | 28.9 (20.5) | 7.9 (0.8) | 17.6 (6.6) |
| Control (n=14) | 13:1 | EA: 13, AA: 1 | 32.7 (16.2) | 8.1 (0.8) | 21.9 (5.2) |
| qPCR Analysis |  |  |  |  |  |
| ASD (n=7) | 5:2 | EA: 6, AA: 1 | 24.9 (17.6) | 6.3 (0.8) | 17.4 (6.7) |
| Control (n=10) | 8:2 | EA: 9, AA: 1 | 33.0 (25.0) | 6.9 (0.9) | 19.6 (4.8) |
| Cytokine/Chemokine Measures |  |  |  |  |  |
| ASD (n=19) | 14:5 | EA: 17, AA: 2 | 30.2 (20.9) | **3.9 (2.0)\*** | 17.3 (6.2) |
| Control (n=9) | 8:1 | EA: 8, AA: 1 | 34.3 (19.7) | **5.8 (1.7)\*** | 20.7 (5.0) |

**Notes:** \* indicates p < 0.05 when comparing ASD versus Controls using a Student's t-test.
**Abbreviations:** AA = African American, ASD = Autism Spectrum Disorder, EA = European American, PMI = post-mortem interval, RIN = RNA Integrity Number, S.D. = standard deviation

### RNA-Seq Analysis

Whole transcriptome RNA-Seq libraries were constructed with the Ion Total RNA-Seq Kit v2 (Thermo Fisher Scientific, Waltham, MA) using RNA isolated from frontopolar cortex of ASD samples and controls and sequenced on the Ion Proton System (Thermo Fisher Scientific, Waltham, MA) at the Cedars Sinai Genomics Core (Los Angeles, CA), as an Ion Torrent Certified Service Provider. We used the resultant FASTQ files to perform pseudo-alignments to human genome build GRCh38 and transcript isoforms from Ensembl Genes release 95 with kallisto v0.44.0 [33], generating transcript abundances across the transcriptome. We subsequently tested for differential expression with sleuth v0.29.0 [34] in R v3.5.1. For additional alternative splicing analyses, we performed full transcriptome alignments with GSNAP v2019-01-24 [35] to human genome build GRCh38. We detected all known and novel splicing events by specifying both the -N tag, to enable detection of novel splicing events, and the -s tag, providing the software with a list of known splice acceptor and donor sites from Ensembl Genes release 95 (Human genes GRCh38.p12). We sorted aligned sequence alignment map (SAM) files and analyzed differentially expressed exons using DEXSeq [36] adjusting for RIN as a covariate. We used the Benjamini-Hochberg procedure to account for transcriptome-wide multiple hypothesis testing, reporting significant genes and exons where the false discovery rate adjusted *p*-value (*q*-value) is <0.05, unless otherwise noted.

### Quantitative PCR

We synthesized cDNA from 500ng of total RNA with SuperScript IV (Invitrogen, Carlsbad, CA), using random hexamers and oligo-dT. We performed quantitative PCR (qPCR) using the QuantStudio 3 System (Applied Biosystems, Foster City, CA), with both custom primers and Power SYBR Green (Applied Biosystems, Foster City, CA) or TaqMan™ probes (Applied Biosystems, Foster City, CA) (Supplemental Table S1) and NEB Luna Universal Probe qPCR Master Mix (New England Biolabs, Ipswich, MA). We measured expression in triplicate in three independent experiments. We used the geometric mean of cycle threshold values (C_T_) for the housekeeping gene *ACTB* as an internal control to normalize the variability, and expression values are reported as C_T_ differences from *ACTB* (Δ*ACTB*). We excluded any sample with RIN < 5 at the time we conducted this experiment from further analysis.

### Protein Expression in Brain

We homogenized brain tissues in a 1x phosphate buffered saline solution with 0.1% Triton X-100 and a protease inhibitor cocktail (Pierce Protease Inhibitor, Thermo Fisher Scientific, Carlsbad, CA) and filtered the supernatant using a 0.22μm spin-based column (Ultrafree-MC, UFC30GV0S, Millipore Corp, St. Charles, MO). We subsequently measured protein levels from filtered supernatants using two approaches. First, we measured 41 different cytokines and chemokines using the Milliplex MAP Human Cytokine/Chemokine Magnetic Bead Panel (Millipore kit no. HCVD3-67CKHCYTOMAG-60K, Millipore Corp, St. Charles, MO). According to the manufacturer’s instructions, we thawed homogenates at room temperature, diluted 1μg of total protein 1:5 with PBS, and incubated in the plate for 12 h at 4 °C. We normalized tissue analyte concentrations to total protein concentrations of homogenate supernatant for each sample, as measured by the Qubit Protein assay (Thermo Fisher Scientific, Carlsbad, CA), and present the data as pg/ml protein homogenate. We excluded analytes where all samples had values below the level of detection, according to their respective standard curves. For the remaining 21 analytes, samples where values were below the level of detection for a given analyte were imputed to the minimal detectable level of that analyte for statistical analyses. The 21 analyzed analytes included FGF-2, FLT3L, Fractalkine, G-CSF, GRO, IFNα2, IL-1α, IL-1β, IL-1RA, IL-6, IL-8, IL-13, IP-10, MCP-1, MIP-1α, PDGF-AA, PDGF-AB/BB, RANTES, TGF-α, TNFα, and VEGF-A. For two additional proteins, PSD-95 and TSPO, we performed standard Western blot analyses, normalizing our proteins of interest to β-actin. We purchased the anti-PSD-95 (catalog #: D27E11) and anti-Actin (catalog #: 13E5) antibodies from Cell Signaling Technology (Danvers, MA) and the anti-TSPO antibody (catalog #: EPR5384) from Abcam (Cambridge, MA).

### qPCR and Protein Statistical Analysis & Visualizations

All statistical analyses and visualizations were performed in R v3.5.1. For qPCR and protein measures, we performed Analysis of Variance (ANOVA) comparing cases and controls using the lm and anova functions in the base stats package in R. We used Akaike Information Criterion (AIC) from the MASS package to select significant covariates for each analysis from age, sex, race, PMI, and RIN. We created boxplot visualizations using the ggplot2 package in R. We transformed outliers, defined as values greater than quartile 3 + 1.5 × interquartile range (IQR) or less than quartile 1 − 1.5 × IQR across the whole range of values in both the control and ASD groups, by capping upper values at the 95^th^ percentile and lower values at the 5^th^ percentile.

### Network and Pathway Analyses

We performed weighted gene co-expression network analysis (WGCNA) using the WGCNA package in R [37] with the top 5000 differentially expressed genes from the RNA-Seq experiment, with a minimum module size of 30. We correlated module eigengenes with demographic factors and protein expression levels, to identify co-expression modules associated with these factors. We subsequently performed ingenuity pathway analysis (IPA) on groups of genes constituting the co-expression networks identified by WGCNA, focusing on networks that significantly correlated with protein expression. For IPA analysis, we included the case *versus* control gene expression fold changes from our RNA-Seq analysis, to detect direction-of-effects for significantly enriched pathways and functions.

### Single Nuclei RNA-Sequencing (snRNA-Seq)

Using publicly available data (Gene Expression Omnibus accession GSE97930; [38]) we performed analyses of gene expression enrichment in different cell types of the human frontopolar cortex. We extracted expression data for the 6,154 cells originating from BA10 and performed cell type clustering with Seurat [39]. We examined the expression of VEGF-A across different cell types. Using AUCell [40] we also tested for the enrichment of differentially expressed genes from our RNA-Seq analysis and for the enrichment of genes associated with VEGF-A from our WGCNA analysis across the different cell types.

## Results & Discussion

### Gene Expression & Protein Analysis Finds Upregulated VEGF-A in ASD

Comparing ASD cases to typically developed controls for our RNA-Seq data, we observed 11 genes that were significantly upregulated in ASD cases (Figure 1). Amongst them was *VEGFA*, encoding the VEGF-A protein, exhibiting approximately a 2-fold increase in ASD frontopolar cortex. We performed qPCR analysis on a subset of genes in the samples with acceptable RNA quality to confirm our RNA-Seq findings. Five of 6 genes we tested with qPCR exhibited higher expression in ASD than controls, consistent with our RNA-Seq findings (Figure 2). Two of the genes trended toward significant differences between cases and controls (*VEGFA*, *p* = 0.058 and *ADGRG1*, *p* = 0.072). When comparing expression measured by RNA-Seq and qPCR for *VEGFA*, we observed significant negative correlation between normalized kallisto gene abundance estimates and *ACTB*-normalized C_T_ values (*r*=−0.62, *p*=0.024), as would be expected, since lower C_T_ values in this analysis represent higher gene expression. The *VEGFA* qPCR primers targeted the exon 2-3 junction, which we found to be more abundant in ASD samples than controls in our RNA-Seq experiment (Supplemental Table S2). Thus, our qPCR findings are consistent with our RNA-Seq results, particularly regarding *VEGFA* RNA expression.

**Fig 1.**
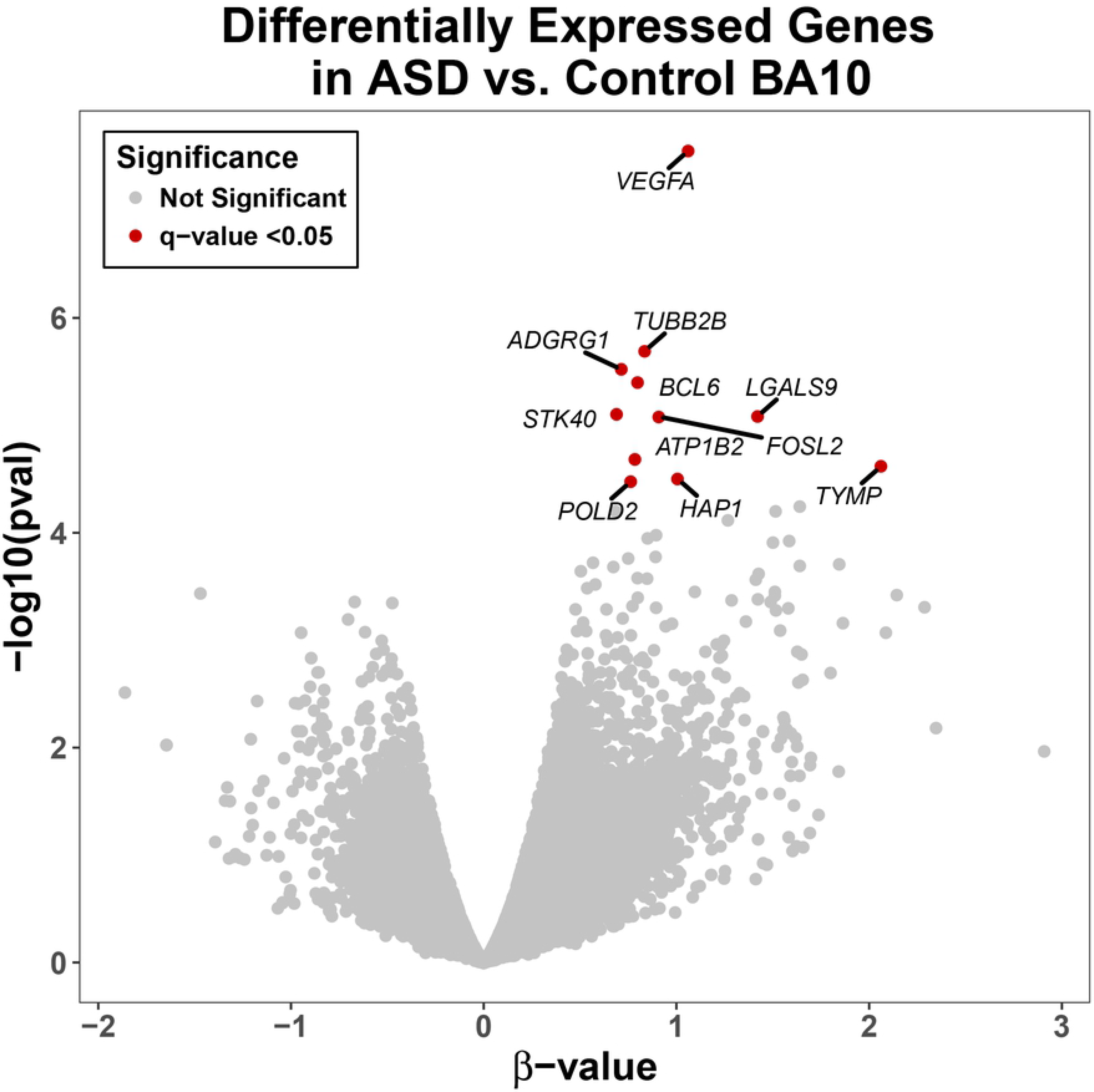
RNA-Seq Volcano Plot. Eleven genes are significantly upregulated in frontopolar cortex brain tissues from individuals on the autism spectrum, according to RNA-Seq analysis. *Note*: A 1-point change in β-value represents approximately a 2-fold difference in gene expression across diagnosis.

**Fig 2.**
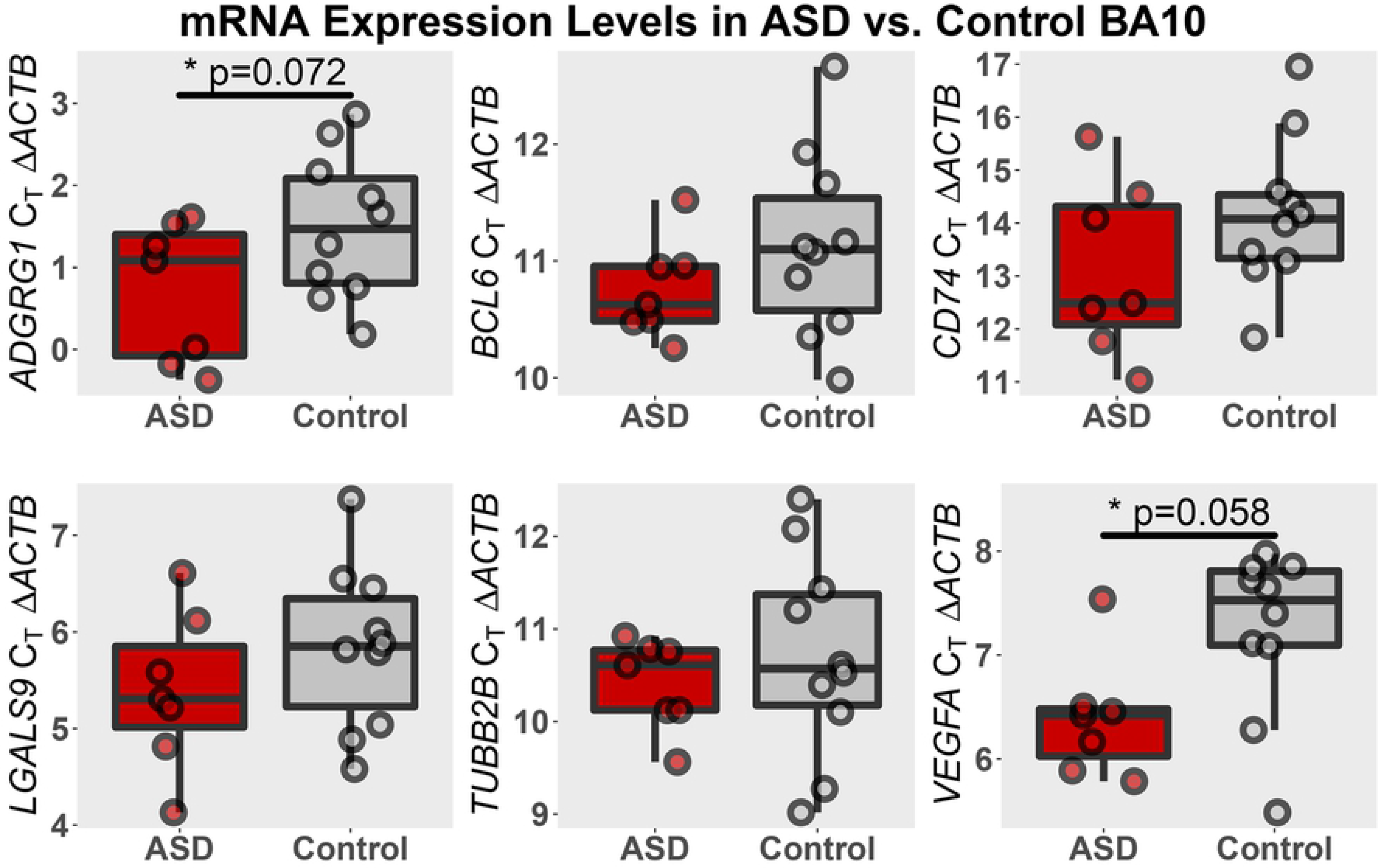
qPCR Validation of RNA-Seq Results. *ADGRG1* and *VEGFA* gene expression in the frontopolar cortex, as measured by qPCR, is consistent with RNA-Seq. ASD samples (*n*=7) express approximately 2-fold more *VEGFA* and 1.5-fold more *ADGRG1* than control samples (*n*=10) and the comparisons for both genes across diagnosis show a trend towards significance. The remaining genes, while consistent with the changes in expression observed via RNA-Seq, were not significant when compared across diagnosis. *Notes*: Gene expression is normalized to β-actin (*ACTB*), with lower numbers indicative of higher expression. The boxplots represent the first quartile, median, and third quartile, while the whisker bars represent the range of values within ± 1.5 × IQR. *Abbreviations*: ASD = Autism Spectrum Disorder, BA10 = Brodmann Area 10, C_T_ = cycle threshold.

VEGF-A plays a complex role in development due to the many different isoforms encoded by the *VEGFA* gene. Based on observations in rodents [41], the developing human brain likely expresses three major VEGF-A isoforms, VEGF-A_121_, VEGF-A_165_, and VEGF-A_189_, with the nomenclature reflecting the size (in amino acids) of the protein. Alternative splicing at exons 6 and 7 create these three isoforms, while alternative splicing of exon 8 confers affinity for the neuropilin-1 receptor (NRP-1) [42,43]. Exon-exon junctions from our RNA-Seq alignments are consistent with the expression of these three VEGF-A isoforms (Supplemental Figure S1, Supplemental Table S2), which include exon 8a – the higher affinity form for NRP-1 binding. We did not find any evidence for usage of alternative exon 8b, associated with reduced neuropilin-1 receptor (NRP-1) affinity [42]. We tested whether any specific isoform was responsible for the increased *VEGFA* expression in ASD by examining differential exon expression transcriptome-wide using DEXSeq. With respect to *VEGFA*, this analysis revealed nominally significant increases in exons 1 and 6a and a nominally significant decrease in exon 8 (Table 2, Supplemental Figure S2). We also found differentially expressed exons from eight other genes that reached transcriptome-wide significance (Table 2). *VEGFA* exons 1 and 8 are constitutive exons for VEGF-A_121_, VEGF-A_165_, and VEGF-A_189_, but exon 6a is only included in VEGF-A_189_. Therefore, we find it plausible that the isoform encoding VEGF-A_189_ is responsible for increased *VEGFA* RNA expression in ASD.

**Table 2.**
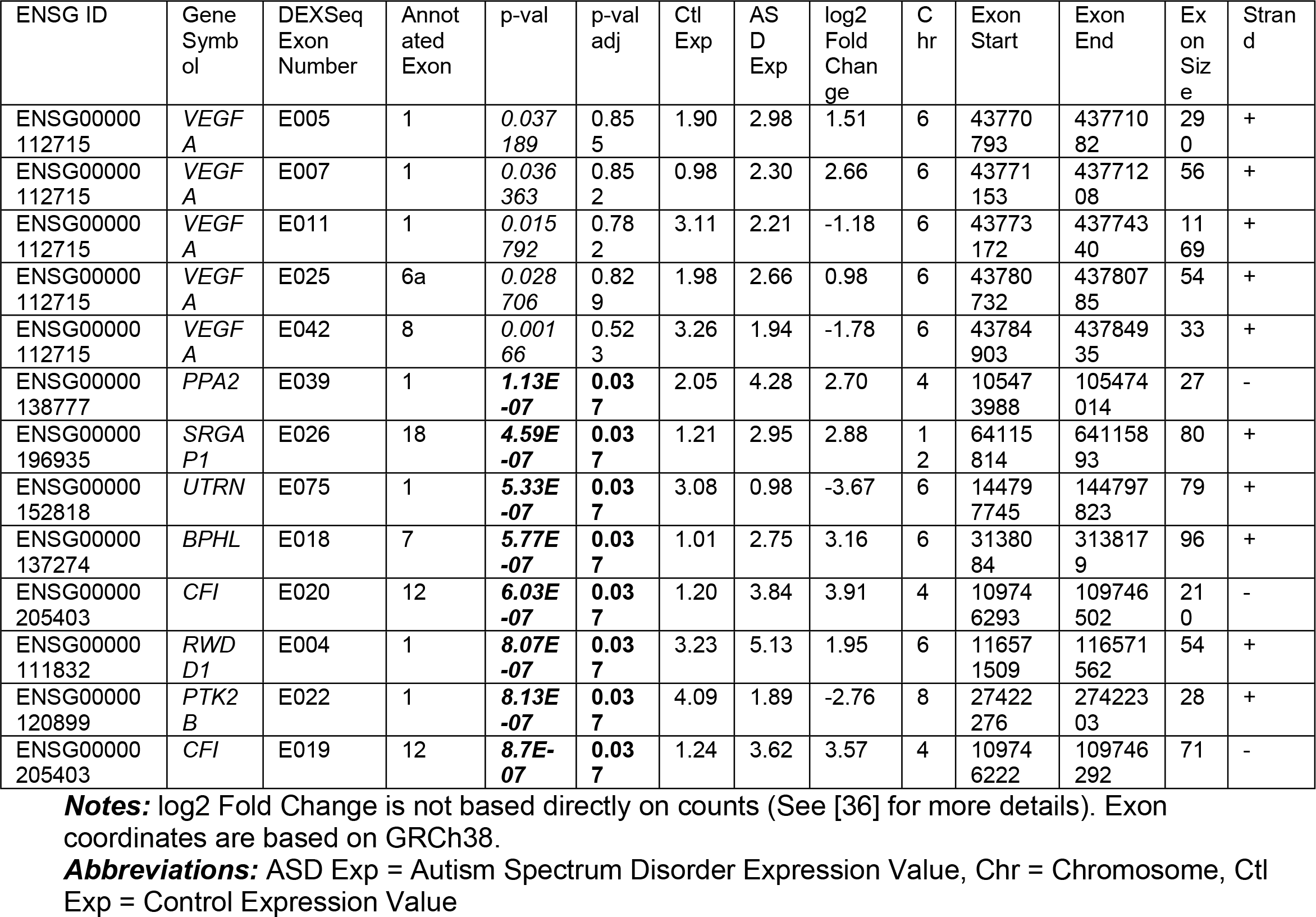
Differentially Expressed Exons from DEXSeq Analysis.

Of the 41 cytokines & chemokines we measured using the Milliplex panel, 21 met acceptable cutoffs for detection and were included in our analyses (Supplemental Table S3). Of the 21 analyzed proteins, IL-1β, RANTES (CCL5), and VEGF-A was significantly increased in the brains of ASD cases versus controls (Figure 3), while GRO (CXCL1), IL-6, and MIP-1α (CCL3) trended towards significant increases (Supplemental Table S3). Here, we note that capping the outliers in our protein analyses resulted in tighter distribution of values, especially across the ASD samples, affecting the significance of the results (Supplemental Table S3). However, the outlier capping did not affect the direction of effects or our interpretation of the data, as the outliers were exclusively ASD samples with greatly increased cytokine and chemokine levels. On average, the ASD cohort expressed approximately 1.4-fold more VEGF-A protein than controls, consistent with the increased RNA expression we observed in the ASD cohort. Western blot analysis to determine the relative expression of each protein isoform was inconclusive, as we could not discern VEGF-A dimers from monomers that use the noncanonical upstream CUG translation initiation site [44]. For example, the VEGF-A_165_ dimer using the AUG initiation site and VEGF-A_189_ monomer using the CUG initiation site are both approximately 44 kDa. Consistent with the protein measures, *CCL5* mRNA expression, encoding RANTES, was higher in ASD samples relative to controls, but this difference was not significant (*p*=0.11, *q*=0.68). *IL1B* mRNA expression, encoding IL-1β, was not detected in our RNA-seq analyses.

**Fig 3.**
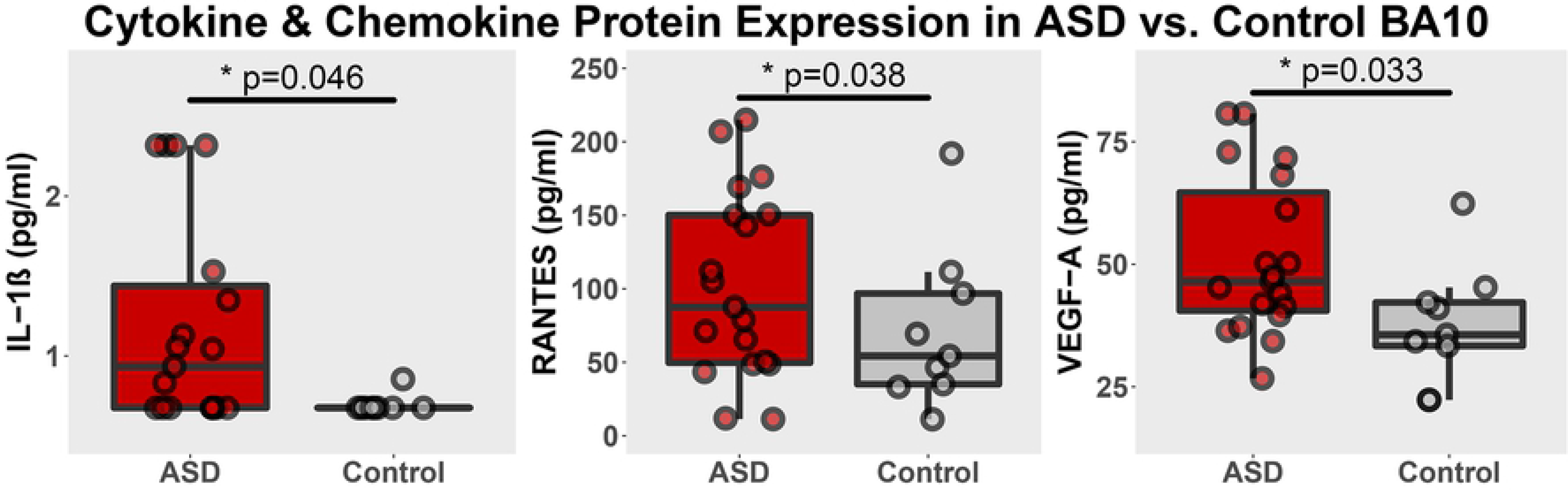
Cytokine & Chemokine Protein Expression from Multiplex ELISA. IL-1β, RANTES, and VEGF-A protein, as measured by the Milliplex MAP Human Cytokine/Chemokine panel, were significantly elevated in the frontopolor cortex of ASD (*n*=19) samples versus Controls (*n*=9). Increased VEGF-A protein expression in ASD is consistent with *VEGFA* RNA expression, as measured by RNA-Seq and qPCR. *Note*: The boxplots represent the first quartile, median, and third quartile, while the whisker bars represent the range of values within ± 1.5 × IQR.

### Network and Pathway Analyses Reveal Dysregulated VEGF-A Signaling and Neuroinflammation

Our WGCNA analysis identified 15 unique co-expression networks. Of particular note, the network eigengene most highly correlated with VEGF-A protein expression (*r*=0.65, *p*=1×10^−4^, arbitrarily named the “green” network; 435 total genes; see Supplemental Table S4 for all gene clusters) is also significantly correlated with 11 additional cytokines and chemokines, including IL-1β (Supplemental Figure S3). Consequently, we focused on the “green” network for pathway analysis using IPA. We performed a Core Analysis using the 435 “green” network genes, revealing multiple lines of evidence that suggests this co-expression network is related to pro-inflammatory signaling. Specifically, the top “Upstream Regulators” are lipopolysaccharide and TNF, the top “Disease and Disorder” annotation is “inflammatory response”, and the top “Physiological System Development and Function” annotation is “immune cell trafficking” (Supplemental Table S5). The Activation Z-scores generated by the gene expression fold-changes consistently finds that the ASD cohort gene expression is associated with greater immune and inflammatory pathways and functions. Additionally, “vegf”, “VEGFA”, “IL1B”, and “CCL5” are all predicted to be significant upstream activators of the “green” gene network (Supplemental Table S5), bolstering the association we observed between this gene network and cytokine and chemokine protein expression. Ingenuity’s VEGFA network, including activation of downstream targets predicted by IPA, are included in Supplemental Figure S4.

To help us discern the role of VEGF-A in the frontopolar cortex, we extracted the frontopolar cortex cells from publicly available snRNA-Seq data [38] and examined gene expression across different cell types. Our clustering analysis found cell populations and cell type-specific markers that were consistent with cortical cell populations in the original publication [38] (Figure 4 and Supplemental Table S6). Following clustering, we examined the enrichment of *VEGFA* and groups of genes, including differentially expressed RNA-Seq genes and those genes included in the “green” WGCNA network, across the different cell types. *VEGFA* was most strongly expressed in oligodendrocyte progenitor cells (OPCs) and astrocytes (Figure 5a, 5b). The group of 11 differentially expressed RNA-Seq genes are enriched in non-neuronal cell populations, including astrocytes, oligodendrocytes, microglia, and OPCs (Figure 6a), while genes from the “green” network are enriched specifically in microglia (Figure 6b).

**Fig 4.**
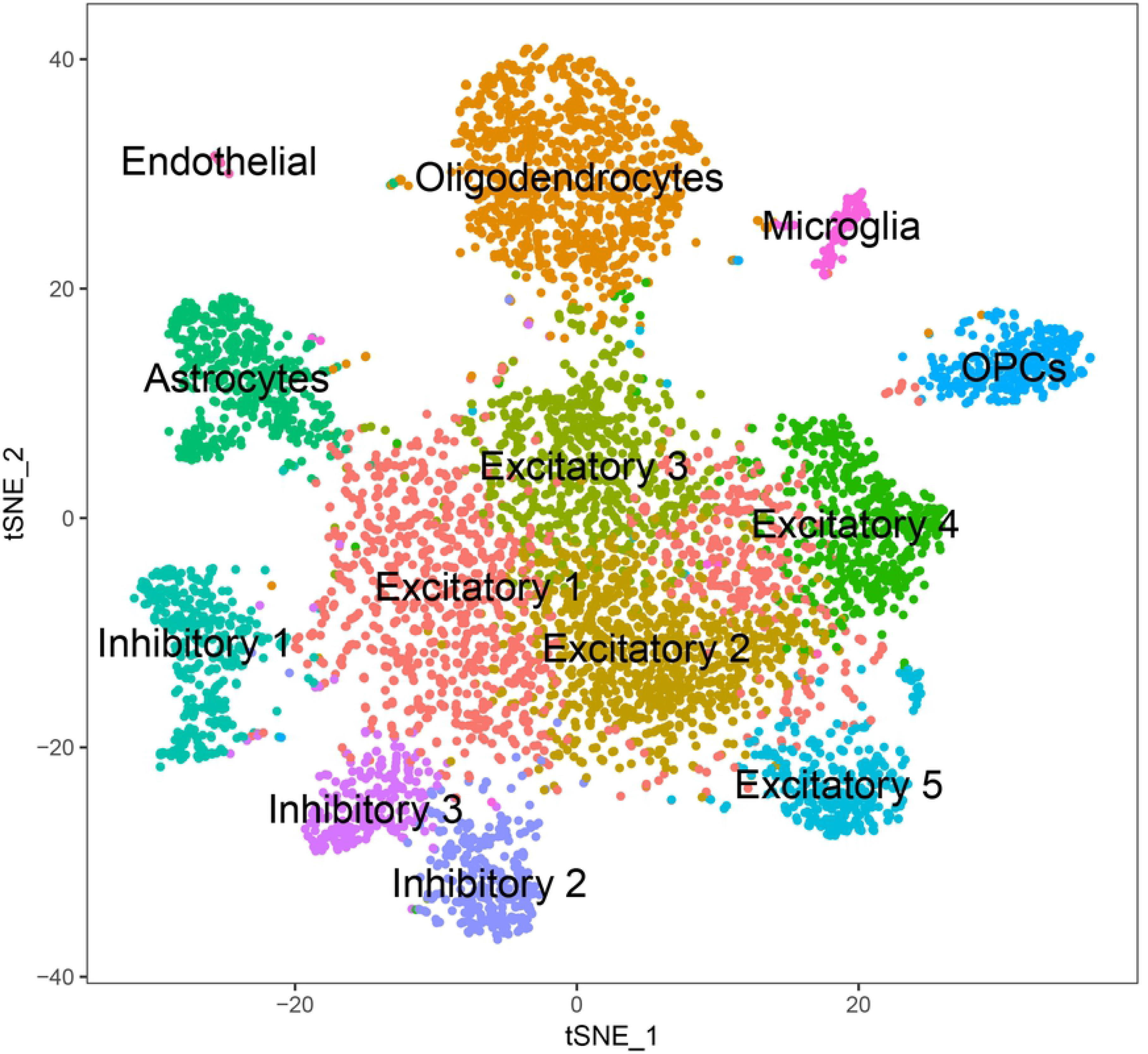
t-SNE Plot of Cell Type Clusters from Single Cell RNA-Seq in Frontopolar Cortex. We clustered 13 different cell types from the frontopolar cortex of typically developed individuals using publicly available data from [38]. The cell type markers we identified are consistent with the markers from the original manuscript (see Supplemental Table S6). *Abbreviations*: OPCs = oligodendrocyte progenitor cells.

**Fig 5.**
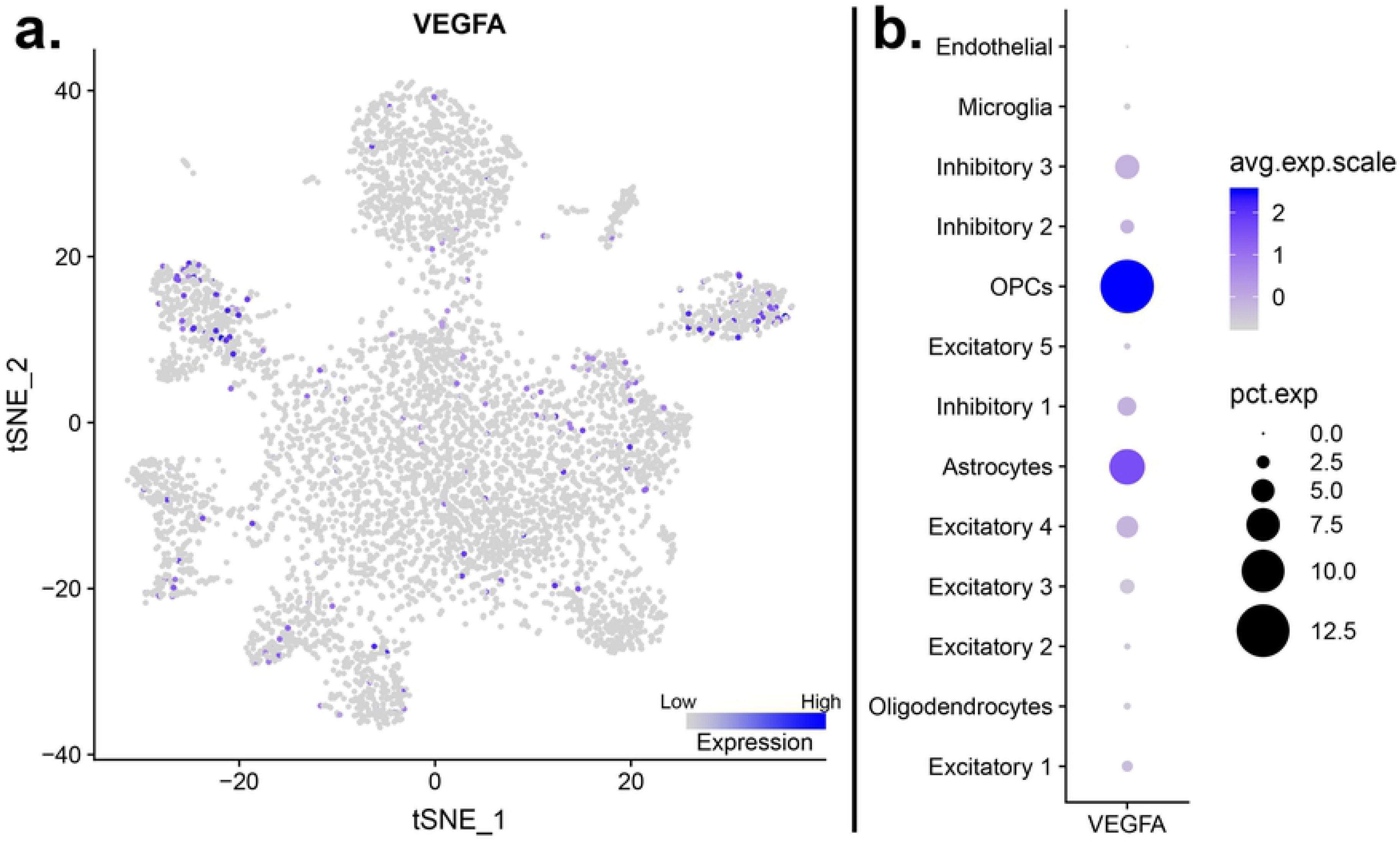
*VEGFA* Expression Across Cell Types in Frontopolar Cortex. a.) The t-SNE plot displaying the expression of *VEGFA* across all of the clustered cells shows sporadic expression of *VEGFA* across most cell types. b.) The dot plot displaying the relative percentage of cells within each cluster expressing *VEGFA* (dot size) and the average intensity of expression within those expressing cells (blue shading) shows that *VEGFA* is most strongly expressed, in terms of number of cells and intensity, in oligodendrocyte progenitor cells (OPCs), followed by astrocytes. *Abbreviations*: OPCs = oligodendrocyte progenitor cells.

**Fig 6.**
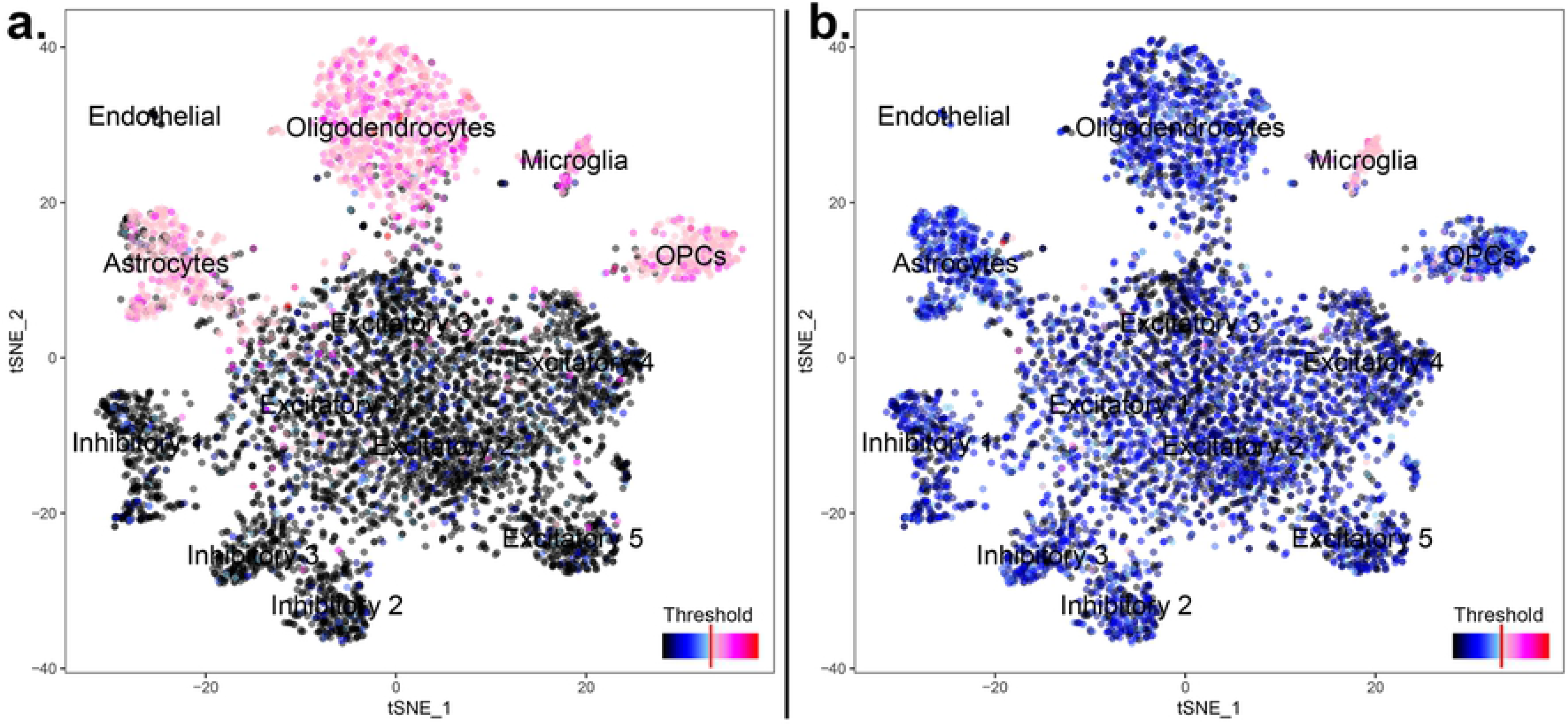
t-SNE Plot of Active Gene Sets Across Cell Types in Frontopolar Cortex. a.) Differentially expressed genes from our RNA-Seq analysis are uniformly enriched in non-neuronal and non-endothelial cell types, as indicated by the pink shading. b.) Expression of genes from the “green” co-expression gene network from our WGCNA analysis is only enriched in microglial cells, as indicated by the pink shading, consistent with this network being associated with neuroinflammation. *Abbreviations*: OPCs = oligodendrocyte progenitor cells.

Although our WGCNA and single-cell analyses find that VEGF-A expression is significantly correlated with groups of genes that are strongly associated with neuroinflammation and enriched in microglia, we cannot discern a cause-and-effect relationship from our analyses. Activated microglia in ASD could stimulate the release of VEGF-A, resulting in a breakdown of the local microvasculature and subsequent inflammation and angiogenesis. Alternatively, increased VEGF-A could be a protective anti-inflammatory response to tonic neuroinflammation observed in ASD. Arguing against a neuroprotective role, the isoform of VEGF-A typically associated with anti-inflammatory response encodes exon 8b [45,46], which we do not see in our splicing analysis.

### Conclusions: Cytokines & VEGF-A in the Context of ASD & Neurodevelopment

Our analyses of RNA and protein expression revealed converging evidence that VEGF-A is increased in the frontal cortex of individuals with ASD, with concurrent increases in the pro-inflammatory cytokines IL-1β and RANTES. Interpreting this finding and understanding its significance requires us to consider the regulation of VEGF-A expression and the many roles VEGF-A plays in normal and aberrant neurodevelopment and ongoing brain-related processes. This includes the diametric roles VEGF-A can play in angiogenesis, neuroinflammation, and neurodevelopment, which depend on VEGF-A isoform expression and the complement of VEGF receptors through which the ligand is signaling. We also must consider peripheral levels of VEGF-A *versus* those observed in the central nervous system, as changes in VEGF-A and its receptors have been detected in the periphery of ASD patients.

Increased IL-1β and RANTES is observed in peripheral blood from ASD patients [47]. Cells from the peripheral immune system of ASD patients show greater IL-1β release in response to their activation [48], perhaps indicative of a subset of patients with a pro-inflammatory phenotype [49]. Related to the gene expression changes we observed, IL-1β was highly correlated with the “green” gene co-expression network, enriched in microglia cells. IL-1β, secreted from glial cells in the CNS [50,51], is a potent stimulator of VEGF-A expression in the context of neuroinflammation and hypoxia [51–53], providing a potential mechanism to explain the increases in VEGF-A we observe in ASD brain tissue. RANTES can also stimulate the release of VEGF from endothelial cells and has pro-angiogenic properties [54]. These findings are consistent with the upregulated levels of VEGF-A we observed in ASD.

Decreased VEGF-A and increased soluble VEGF receptor 1 was observed in the serum of severely affected patients [29], while decreased plasma VEGF-A levels were associated with increased severity of ASD presentation in females [31]. These findings in the periphery differ from our results, as we found that increased VEGF-A was associated with ASD. Peripheral measures of VEGF-A expression do not reflect central measures [55,56], even in patients with brain tumors that express exponentially more VEGF-A [57–59]. This lack of correlation strongly suggests that we cannot infer VEGF-A concentration in the CNS based on peripheral measures. However, elevated VEGF-A in the brain can be detected in the CSF [57]. In support of our findings, Vargas et al. [32] found an 81.8-fold increase in VEGF-A in the CSF of ASD patients versus controls. In this study, however, the ASD cohort was significantly younger than the control population (ASD age range 3-10 years old; Control age range 12-45 years old), which is a major confound given VEGF-A’s role in neurodevelopment. Our cohorts did not significantly differ by age (*p*=0.6), nor did VEGF-A levels correlate with age, excluding age as a confounding factor. Since our results are specific to the frontal cortex and not the periphery, it is critical for us to consider the isoform(s) of VEGF-A expressed here, their most likely cell type of origin, and receptor expression throughout neurodevelopment.

Our RNA-Seq analysis did not reveal differential expression for any of the genes encoding canonical VEGF receptors (*FLT1*, *KDR*, and *FLT4*), neuropilin receptors (*NRP1* and *NRP2*) or VEGF-B (*VEGFB*), allowing us to focus on the VEGF-A ligand. Furthermore, we did not see significant increases in RNA expression for *IL1B* or *CCL5*. The developmental effects of increased VEGF-A expression in the brain varies, depending on which VEGF-A isoform is upregulated. For example, overexpression of mouse Vegf-A_120_ (analogous to human VEGF-A_121_) in neurons increases cell density and angiogenesis in the forebrain and appears to have anxiolytic and antidepressant effects on behavior [60]. However, the neurogenic effects dissipates by adulthood [60], suggesting that earlier developmental periods are more responsive to changes in VEGF-A levels. Overexpressing human VEGF-A_165_ in mouse neurons significantly increases capillary density in the frontal cortex and increases blood flow in white matter under severe hypercapnia [61]. Overexpressing VEGF-A_165_ also increases blood-brain barrier (BBB) permeability [62]. Consistent with the consequences of VEGF-A overexpression observed in rodents, ASD patients have increased neurogenesis in the frontal cortex [63,64], persistent angiogenesis [65], increased white matter metabolic rates in the frontal lobes [66], and apparent altered BBB permeability [67,68].

Few studies currently link VEGF-A dysregulation to ASD, perhaps owing the lack of published studies that have examined this growth factor in ASD brain tissues or etiologic heterogeneity across ASD patient populations. Given this caveat, we cautiously interpret our findings and encourage replication studies and those that examine VEGF-A levels in the brains of ASD patients *in vivo*. For example, the anti-VEGF antibody bevacizumab has been radiolabeled with zirconium-89 (^89^Zr-bevacizumab) and used in clinical applications, showing some uptake into the brain [69,70]. However, the utility of ^89^Zr-bevacizumab as an imaging marker capable of detecting modest changes in VEGF-A levels in the central nervous system is yet to be demonstrated.

Even with cautious interpretation, there are three observations worth emphasizing. 1.) Increased VEGF-A can create anatomic and physiological features that are commonly observed in ASD. 2.) Increased VEGF-A can indicate ongoing pathophysiology related to neuroinflammation. 3.) Glial-cell mediated neuroinflammation influences synaptic development and neurogenesis. The question remains whether these observations converge in ASD and account for a significant portion of ASD pathology.

## Acknowledgements

The authors sincerely thank the families of the tissue donors, the Autism Tissue Program, and Autism Brain Net for providing the tissues used in this study. We also acknowledge the assistance of Wolfgang Sadee’s laboratory in generating the RNA-Seq data. This research was supported by a Developmental Start-Up Grant from the University of Iowa College of Pharmacy. *Gene expression files are under submission to Gene Expression Omnibus (GEO). We will update this section with the Accession number once available.*

## Author Contributions

**A.A.:**Investigation, Formal Analysis, Supervision, Writing – Original Draft Preparation, Writing – Review & Editing

**M.N.K.:**Investigation, Writing – Review & Editing

**N.K.B.:**Resources, Writing – Review & Editing

**R.M.S.:**Conceptualization, Data Curation, Formal Analysis, Funding Acquisition, Investigation, Methodology, Project Administration, Resources, Supervision, Visualization, Writing – Original Draft Preparation, Writing – Review & Editing.

## List of Supplemental Tables & Figures

Supplemental Table S1. PCR Primers and TaqMan^TM^ Probes for qPCR Analyses

Supplemental Table S2. *VEGFA* Exon-Exon Junction Counts from RNA-Seq

Supplemental Table S3. Analysis Results of 21 Cytokines/Chemokines from Milliplex Panel

Supplemental Table S4. WGCNA Network Results and Associations with VEGF-A

Supplemental Table S5. Top Findings from Ingenuity Pathway Analysis into the “green” WGCNA Network

Supplemental Table S6. Cell Type Markers Comparing the Current Study and Lake et al. [38]

Supplemental Figure S1. Sashimi Plot for *VEGFA* Splicing

Supplemental Figure S2. DEXSeq Plot Comparing Exon Expression for *VEGFA* in Cases and Controls

Supplemental Figure S3. Correlations between WGCNA Module Eigengenes and Phenotypes

Supplemental Figure S4. VEGF Network from Ingenuity Pathway Analysis

